# Robust discovery of causal gene networks via measurement error estimation and correction

**DOI:** 10.1101/2023.05.09.540002

**Authors:** Rahul Biswas, V P Brintha, Amol Dumrewal, Manikandan Narayanan

## Abstract

Discovering causal relations among genes from observational data is a fundamental problem in systems biology, especially in humans where direct gene interventions or perturbations are unethical/infeasible. Furthermore, causality is emerging as an integral factor for building interpretable and generalizable machine-learning models of complex phenotypes. Existing methods can discover causal relations from observed gene expression and matched genetic data using the well-established framework of Mendelian Randomization. But, the prevalence of expression measurement errors can mislead most existing methods into making wrong causal discoveries, especially among genes transcribed at low to moderate levels and using data with large sample size (say thousands as in modern genomic or GWAS studies).

In this study, we propose a new framework for causal discovery that is robust against measurement noise by extending an established statistical approach CIT (Causal Inference Test). We specifically developed a two-stage approach called RCD (Robust Causal Discovery), wherein we first estimate measurement error from gene expression data and then incorporate it to get consistent parameter estimates that could be used with appropriately extended statistical tests of correlation or mediation done in the original CIT. By quantifying and accounting for noise in the data, our RCD method is able to significantly outperform the baseline method in recovering ground-truth causal relations among simulated noisy genes and transcription factor to target gene relations among noisy yeast genes using data on 1012 yeast segregants. Encouraged by these results, we applied our RCD to a human setting where perturbations are infeasible and identified several causal relations, including ones involving transcriptional regulators in the skeletal muscle tissue.

**Data and Code Availability:** The code that implements our two-stage RCD framework is available here: https://github.com/BIRDSgroup/RCD; code for reproducing the figures/tables in this manuscript is also provided in this link.

## Introduction

Deciphering the genotype→phenotype map and its underlying cause-effect relations has been a longstanding goal of systems biology ([1]), and very recently also a key step in realizing machine learning models that can use causal information to make interpretable and generalizable predictions of disease endpoints ([2]). Discovering the causal network among genes and disease traits is a challenging endeavor. Established means of causal inference using perturbation/knockout experiments or randomized controlled trials are infeasible or unethical in *in vivo* settings like human studies, and it becomes necessary to use observational data alone to learn causality ([3]). In this regard, analysis of data from observational studies such as GWAS (Genome-Wide Association Study) or eQTL (expression Quantitative Trait Loci) studies have revealed not only genetic variants associated with disease and gene expression traits but also gene regulatory networks ([4–8]) and causal mediators of clinical/disease endpoints ([9–11]). Many of the established gene regulatory network discovery methods are based on Mendelian Randomization (MR, [12]), which is a framework that uses a genetic variant as an instrumental variable to test for a causal relationship between two other trait variables (e.g., two gene expression traits) using mediation/conditional-independence or other similar tests. Another well-established method CIT (Causal Inference Test ([5])) uses a statistical testing framework that is more similar to the Baron and Kenny framework ([13]) than the MR framework, but its goal is similar to other MR-based methods, which is to discover causal relations and provide a score or p-value that quantifies the strength or uncertainty of the inferred causality.

A key aspect of observational data often underlooked in current causal discovery studies is measurement errors, despite the prevalence of such errors in high-throughput data ([14–16]) and convincing evidence from a few studies on the deleterious impact of these errors on causal discovery ([1, 7, 17]). Some alternatives have been suggested to tackle this issue ([17]), but mitigating the harmful effects of noise on causal calls remains an important open problem, especially when dealing with noisy genes and datasets of large sample sizes (as is the case with modern GWAS or other genomics datasets). To elaborate, it is well-known that measurement noise is prevalent in gene expression data, like in the integer gene counts measured via the RNA sequencing (RNAseq) technology; and the error magnitude is different for different genes with low to moderately expressed genes typically more noisy than highly expressed genes ([14–16]). These errors, also known as technical variability or noise, could arise from different sources ([18]) like random sub-sampling steps involved in library preparation or sequencing, and bias due to read-mapping ambiguity. Noisy measurements of genes can result in inconsistent or attenuated estimates of the parameters of a linear regression model relating multiple genes ([19]). Since many MR-based or other causal discovery methods rely on parameter estimates of linear regression models, this would mean a loss of power for detecting causal relations at best (or) reversal of the causal direction at worst in the presence of measurement noise ([1]).

While almost all differential gene expression methods acknowledge the prevalence of technical noise in expression data and adopt the best practice of accounting for this noise in their analysis to derive reliable findings (e.g., [20],[18]), only a few gene network discovery studies have assessed the impact of measurement errors on inferring gene coexpression networks ([21, 22]) or gene regulatory networks ([7, 17, 19]). It is high time that this issue is addressed adequately to enable reliable discovery of the causal networks underlying the genotype→phenotype map.

In this work, we propose a two-stage framework for causal discovery that is robust against measurement error by extending the well-established CIT method ([5]) mentioned above. We call our newly proposed method RCD for Robust Causal Discovery – RCD’s first stage estimates the magnitude (variance) of measurement errors when gene expression is quantified using RNAseq, and the second stage uses these error variances to correct the relevant statistics, parameters, and p-values of CIT’s four regression-based statistical tests, which verify a chain of conditions of causality. Our method RCD shows increased statistical power than the baseline method CIT in both simulated data and real-world 1012 yeast segregants’ data, yielding in general more causal calls among noisy genes, at similar false-positive rates. Furthermore, RCD was able to discover an *in vivo* human gene regulatory network operating in the skeletal muscle tissue, comprising known and novel causal relations.

## Results

### Our RCD method overview

RCD takes genetic and gene expression data from the same set of individuals as input and infers causal relations among gene expression variables, one pair at a time. If *G, T* are a pair of genes to be tested for causal relationship and *L* is a SNP associated with both *G* and *T*, then RCD takes such a query trio (*L, G, T*) as three input vectors (see Figure 1), estimates the measurement error of *G* and *T*, and performs a chain of statistical tests by incorporating the measurement errors. Please see Figure 1 for an overview of RCD. In a bit more detail, RCD works in the following two stages to be robust against measurement errors.

**Fig 1.**
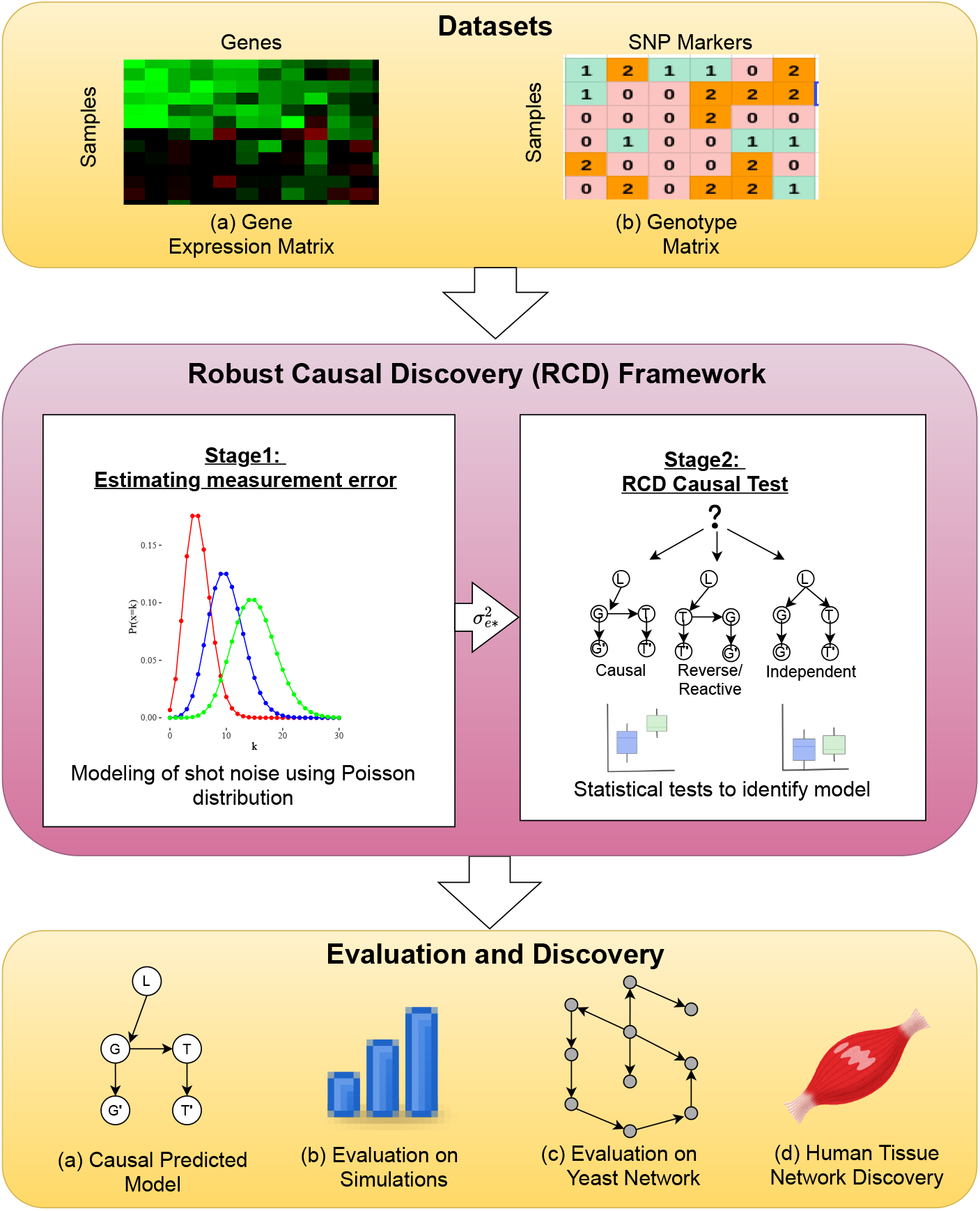
RCD (Robust Causal Discovery) Study Overview. Gene expression and genetic data observed across a set of individuals are analyzed, one trio at a time, using our error-aware two-stage RCD method; and the resultant causal relations are collated and assessed under different simulation/real-world application scenarios. For each trio (*L, G, T*) observed across a set of individuals, RCD analyzes the input genotype vector *L* and the expression vectors of two genes associated with *L* (with *G*′ and *T* ′ indicating noisy measurements of the true unobserved gene counts *G* and *T*). RCD works in two stages as shown to infer the causal relation between *G* and *T* using *L* as an instrumental variable. Please see text for more details.

Stage 1: Measurement Error Estimation: For each gene, this stage uses the gene’s expression data to estimate the magnitude/variance of measurement error of the gene (denoted 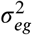 for *G* and 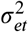 for *T*). RCD can work with different error estimation techniques (see Methods); however our main contribution in Stage 1 is to propose an error estimation technique that works for any gene whose expression is quantified via RNAseq read counts, specifically by modeling sampling noise as a Poisson distribution. One challenge here is to estimate error variance in the normalized gene count space, which we address by sampling dummy integer gene counts from a Poisson distribution with average expression matching the actual gene counts, and performing standard normalization and transformation before computing its variance.

Stage 2: RCD Causal Test: This stage uses the observed data (*L, G*′, *T* ′) and the two gene’s error estimates (denoted 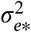) from Stage 1 as input to perform statistical tests of causality between genes *G* and under certain assumptions about the trio. Our main contribution in Stage 2 is to extend*T*the statistical tests of CIT to incorporate measurement error estimates, and more specifically to infer error-corrected estimates of regression model coefficients, residuals, and p-values associated with these tests (thereby making the tests more consistent and robust against noise).

Please refer Methods for a complete description of our error-aware, two-stage RCD method.

### Robustness of RCD on Simulated Data

Methods such as CIT use conditional independence tests as one of the component statistical tests. Previous studies have shown that the presence of measurement errors in gene expression data can lead to the failure of conditional independence tests ([1, 23]). Such failures make causal discovery approaches like CIT suffer from high false negatives, which worsen as the sample size increases ([17]). Our RCD method uses a handle on noise magnitude to make conditional independence tests more robust against noise. To verify if this is indeed the case, we generated simulated data according to a variety of parameter configurations (648 in number; see Methods section on “Simulation Setup”). Across a subset of these configurations, a basic power (True Positive Rate) and type 1 error (False Positive Rate) comparison of CIT and RCD is shown in Figure 2. The level of noise *<*_*et*_ in the outcome variable is fixed, and the level of noise *<*_*eg*_ in the mediator is varied in Figure 2(A). As the level of noise increases in the mediator, the power of CIT decreases rapidly than RCD across all sample sizes. For CIT, the power (TPR) decreases as the sample size increases. In contrast, our RCD performs better with large sample sizes and its power is almost equal to the configuration when measurement noise is zero, thus validating our strategy of incorporating error information for reliable causal discovery. Increased power at similar false-positive rates is also observed when the model is independent (*G* ← *L* → *T*) as shown in Supplementary Figure S1.

**Fig 2.**
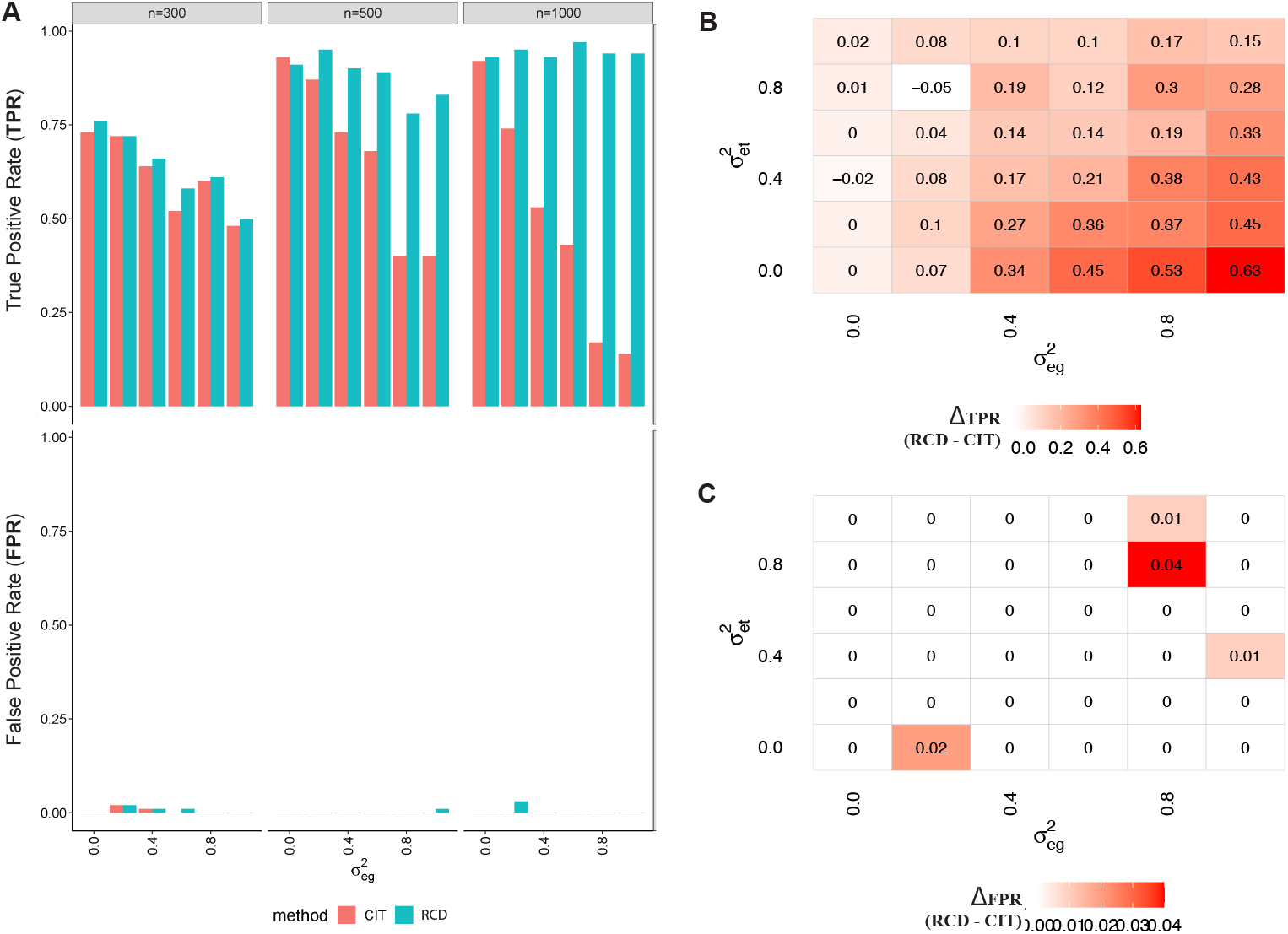
Performance comparisons of our method RCD on simulated data. (A) The top graph plots the true positive rate (fraction of all true causal relations that are inferred as causal by RCD or the baseline method CIT), when data pertaining to 100 true causal pairs are simulated according to the causal model *L* → *G* → *T* (with correlation coefficient 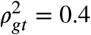). The bottom graph plots the false positive rate (fraction of all non-causal/independent pairs inferred as causal by a method), when data on 100 independent pairs are simulated using the model *G* ← *L* → *T*. Measurement error in trait *G* 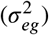 is varied along the x-axis keeping the measurement error in trait *T* fixed at 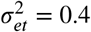. The power (TPR) of CIT decreases steeply with increasing noise in *G*, whereas RCD performs almost the same as on error-free data (for higher sample sizes; columns show different sample sizes, *n* = 300, 500, 1000). (B, C) The effect of varying measurement errors in both *G, T* is shown (for *n* = 500). RCD has better TPR than CIT when measurement error in *G* is higher than that of *T* at similar FPR rates.

To inspect RCD’s performance more extensively, we also allow the error magnitude of *T* to vary from 0 to 1, and the resulting performance of RCD relative to CIT is shown in Figure 2(B,C). For almost all configurations, the difference of power between RCD and CIT (Δ_*TPR*_ = *RCD*_*TPR*_ − *CIT*_*TPR*_) is positive and also larger for higher levels of noise in *G* (Figure 2(B)). Figure 2(C) shows that RCD is not compromising in terms of false positive rates also.

### RCD’s Recovery of GRN among Noisy Yeast Genes

To compare the performance of RCD with CIT on a real-world network, we applied both the methods on a ground-truth yeast causal network (DNA binding plus Expression network from Yeastract ([24]). This ground-truth network captures transcription factors (TF) and regulated target genes (TG) experimentally validated through TF-DNA binding interactions or differential expression upon perturbation of TF. To verify that RCD can recover causal relations among highly noisy genes, we used two subsets of the ground-truth networks (i) On the overall trios, both methods are better than a random classifier, with RCD having AUPR_k%_ better than CIT for various values of *k*%. (ii) On trios whose technical to total variance ratio is high (≥ 0.4; see Figure 3(B)), specifically when 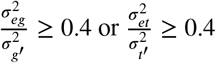, the performance of CIT on these highly noisy trios drops because it results in more false negatives and at some point becomes worse than a random classifier, whereas RCD is performing better than CIT and the random classifier (Figure 3(C)). For both the methods, identifying the target trios filtered on highly noisy genes is more challenging than the overall trios. The gap between CIT and RCD on the filtered trios is much wider and hence shows the robustness of RCD in the presence of noise and potentially a better causal mediation test in such real-world settings. When trying two other technical to total variance cutoffs, 0.3 and 0.5, similar performance trend of RCD outperforming CIT is observed for the latter high-noise cutoff (Supplementary Figure S2).

**Fig 3.**
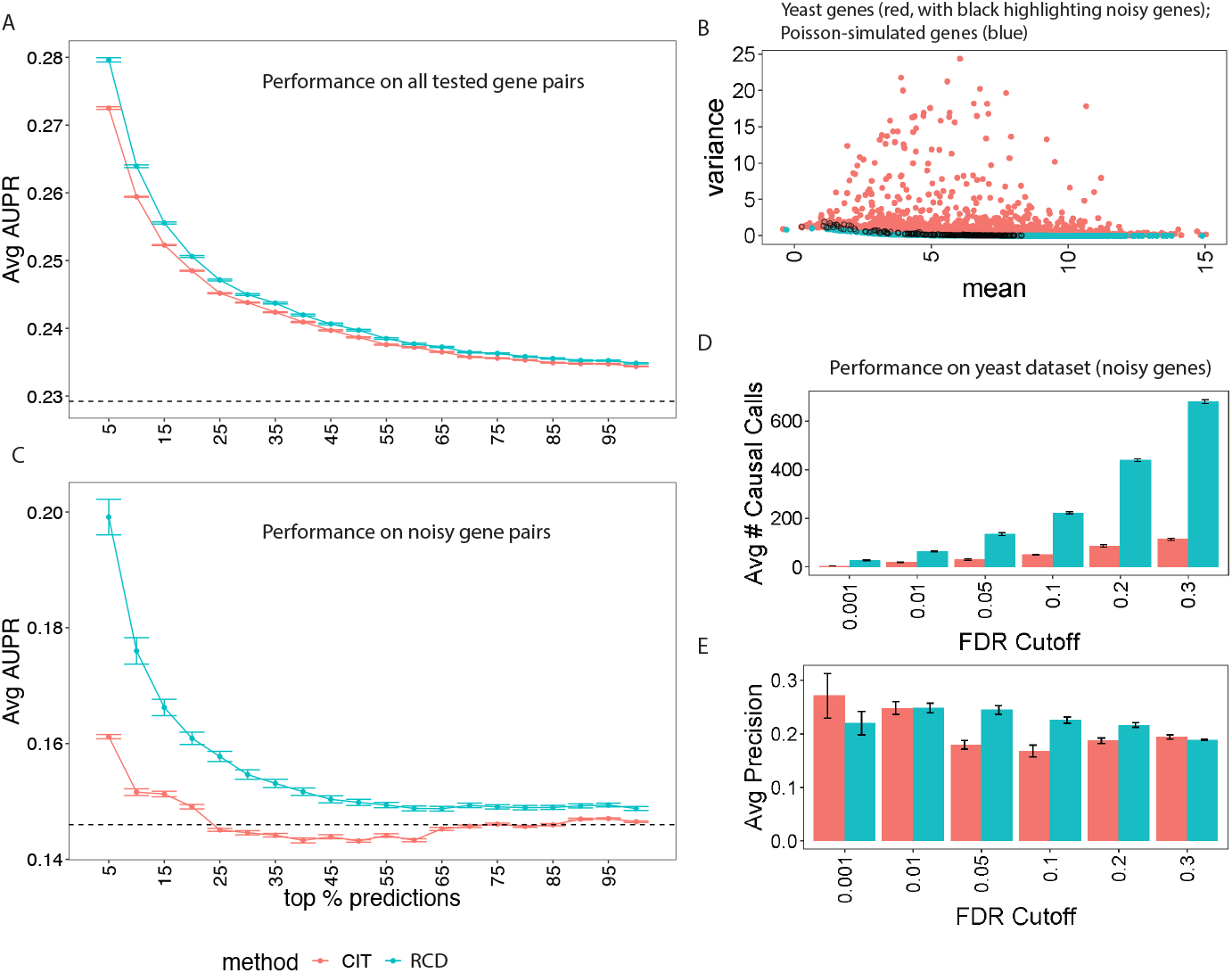
Performance of our RCD relative to the baseline CIT on yeast dataset. The goal is to recover ground-truth causal regulation matrix of TFs→TGs. From the ground-truth regulation matrix, these two query trios sub-lists are constructed and tested separately: (A) a complete query list of trios having 62052 causal relations and 207933 non-associations, and (B) a noisy subset comprising only trios with high measurement errors (see Methods; specifically trios with genes whose error variance is at least 40% of the total variance (C), yielding 4773 causal relations and 28022 non-associations). (A, C) Among the *top*_*k*%_ causal pairs (selected based on CIT or RCD causality p-values), average AUPR is computed. The average performance across four runs for different values of *k* is shown, with error bars indicating standard deviation across these runs. (D, E) We also show the performance of RCD vs. CIT using standard Benjamini-Hochberg adjusted p-values (FDR cutoff in x-axis). At different FDR cutoffs, relative to CIT, our method RCD gives more causal calls (D) at comparable or better precision (E).

The better performance of RCD over CIT in a real-world setting is promising as it validates our overall framework comprising both Stage 1 error estimates and Stage 2 correction procedures. Many of our model assumptions like linear causal relations among genes and independent additive Gaussian errors could be viewed as very simplistic representations of a real-world dataset, still we can see tangible benefits among noisy genes in terms of the relative performance of RCD over CIT. Another way to see the better performance of RCD is to observe how many calls each method makes at different FDR cutoffs (i.e. Benjamini-Hochberg adjusted p-values that account for multiple testing) and the precision of these causal calls (Figure 3(D,E)).

### RCD’s Inferred Transcriptional Regulators in Human Muscle

Gene regulatory networks encode the complexity of the biological system. In humans, we have about 20,000 genes and hence roughly 200 million gene pairs to consider to test for causality. With such a large search space, conducting gene perturbation or intervention experiments is infeasible, and on top of that, the unknown consequences of such perturbation would make it unethical. Hence in such a real-world human setting, using observational data to identify novel putative causal gene pairs will be of great value.

We would like to assess the value of RCD in such a human setting. To do so, we applied RCD to the human skeletal muscle tissue of the NIH GTEx (National Institutes of Health, Genotype-Tissue Expression [25]) data. RCD was able to uncover a gene regulatory network having 314 genes and 164 edges, and the degree properties of this whole human-muscle-specific network is summarized in Table 1.

**Table 1.** Out-degree property of the inferred human network. The human skeletal muscle gene regulatory network discovered by RCD has 314 genes and 164 directed edges. Direction *G* → *T* is predicted if p-value_*RCD,G*→*T*_ < 0.05 and p-value_*RCD,T* →*G*_ > 0.05. Of the 157 putative regulators identified, 153 are predicted to regulate one gene.

| Number of putative regulators | Number of target-genes (Out-degree) |
| --- | --- |
| 1 | 5 |
| 3 | 2 |
| 153 | 1 |

A part of the whole network having module (connected component) sizes of more than 2 is shown in Figure 4(A). The network inferred by the RCD in Figure 4(A), identified KLF5 as a potential hub regulator driving 5 target genes in human skeletal muscle tissue. Previous studies have experimentally shown the pivotal role of KLF5 in the skeletal muscle tissue of mice ([26]). In brief, they have shown essential roles of KLF5 such as deletion of KLF5 impairs muscle regeneration after injury, knockdown of KLF5 suppresses differentiation of myogenesis process, and inhibition of KLF5 affects transcription regulation of muscle-related genes. For instance, Figure 4(B) shows one such example of SNP 13_74110412→KLF5→PHETA1 causal relation identified by RCD, where KLF5 is regulating the target gene PHETA1 and the SNP’s effect on PHETA1 vanishes when conditioned on KLF5. We also show an example gene pair in Supplementary Figure S4 where there is lack of evidence for such a vanishing of the SNP’s effect, and therefore RCD calls the gene pair as spuriously coexpressed due to the shared confounding SNP. These results taken together, in a complex human setting where perturbation experiments are infeasible, show that RCD offers a route to reliable causal discovery and identifies tissue-relevant TFs.

**Fig 4.**
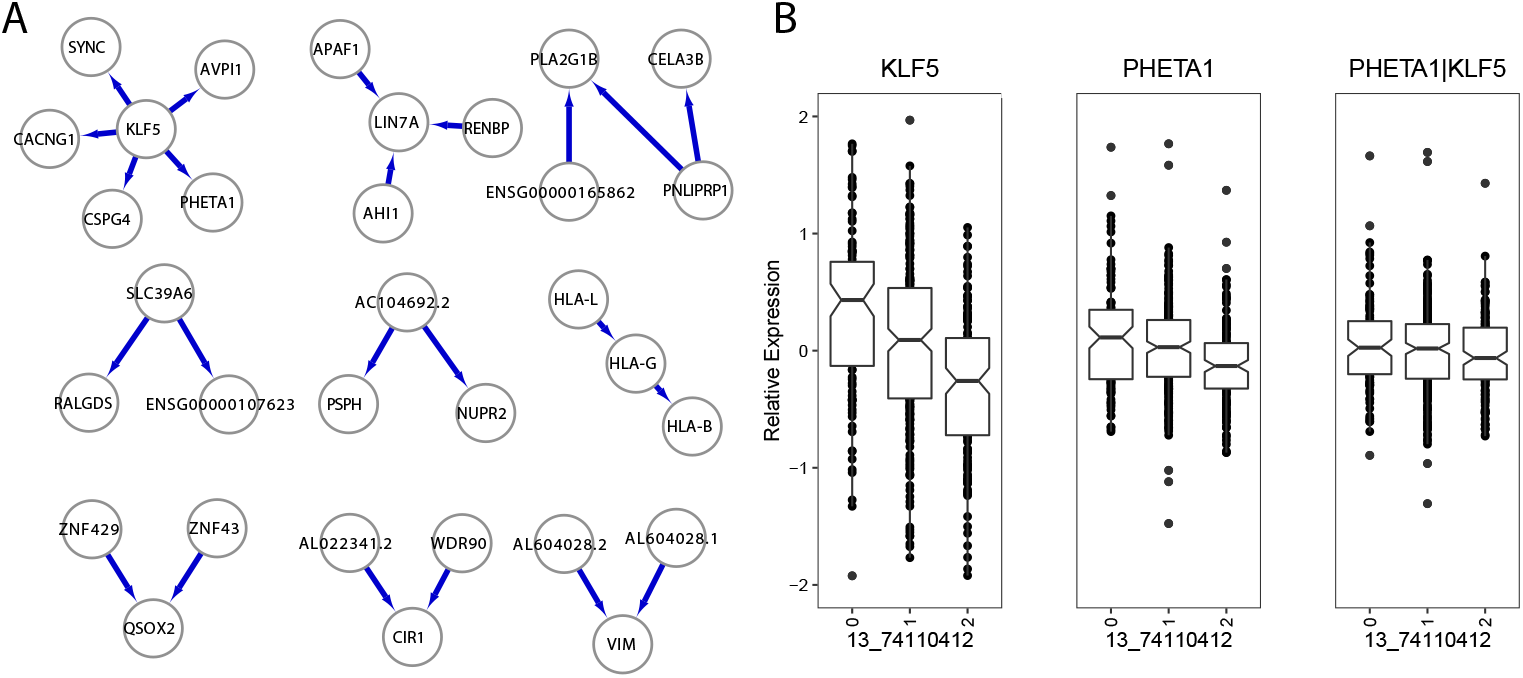
Gene regulatory network in human skeletal muscle identified by RCD. (A) Only connected components of this gene network having more than 2 genes are shown (see Supplementary Figure S3 for the full network). Network shows KLF5 as one of the potential transcription factors in the skeletal muscle tissue regulating many genes. (B) An illustration of causal mediation using a potential transcription factor KLF5 identified in skeletal muscle. A SNP 13_74110412 (*L*) is used as the instrument (encoded as 0, 1 or 2, based on the copy number of alternate allele) to detect causal relation *L* → *G* (KLF5) → *T* (PHETA1). The causal relation is evident from the effect of the SNP on *G* and *T* in the first two plots, and the vanishing of this effect on PHETA1 once conditioned on the mediating regulator KLF5.

## Discussion

Measurement error is prevalent in gene expression data and can mislead causal gene network discovery methods into making wrong or inconsistent causal inferences ([7]). In this study, we have proposed a statistical approach for causal discovery to be robust against measurement noise, with a specific focus on extending the well-established CIT method. Our two-stage RCD method quantifies and accounts for measurement errors in the data. By doing so, RCD showed power improvements over CIT across different levels of measurement errors in simulation studies, and especially for genes with moderate-to-high error levels and at large sample sizes. When applied to yeast data comprising 1000+ segregants, RCD was able to recover causal relations among noisy genes better than CIT. Our method, when applied to the muscle tissue of the NIH GTEx data ([25]), revealed transcriptional regulators in muscle such as KLF5 and led us to build an *in vivo* human gene regulatory network.

There are certain caveats with any MR-based causal discovery method in general and our RCD method in particular that is worth mentioning. All MR-based methods can typically recover only a small fraction of all ground-truth causal gene interactions from observational data, since not all causal gene pairs will have a shared associated genetic variant *L*, not all trios will satisfy MR assumptions, and samples sizes may be insufficient to detect weak causality. Nevertheless, the significant causal relations detected by MR-based methods or RCD are reliable, and could potentially reveal hundreds of novel gene regulatory interactions, which can then be probed experimentally. Regarding RCD-specific caveats, our gene-specific error estimates are average error magnitude across all samples, and not sample-specific for simplicity. By focusing on the CIT framework in RCD, we also inherit some of the assumptions of CIT such as Gaussian distribution for continuous variables, and linear relationships among the variables. It would be interesting to see how non-linear causal relationships can be learnt under different distributional assumptions (such as Negative Binomial) robustly in the presence of measurement noise.

Despite the simplifying model assumptions adopted by our RCD method, we show that incorporating error information has clear benefits for causal discovery from both simulated and real-world genomic datasets. We can foresee reaping more benefits with richer models of gene expression data and causal relations among genes.

The measurement error modelling we employ in our current study is based on the mean-variance relationship of a Poisson model for estimating technical noise. We could also employ machine learning methods that use features other than average (mean) gene expression, such as gene length, GC content, etc., to refine our predictions of measurement noise in the future. Quantifying uncertainty in gene expression data can be from multiple sources such as measurement error due to random sampling involved in library preparation steps (shot noise) or due to sequencing bias or due to mapping ambiguity or any others factors. In a recent study, the decomposition of variance of the gene into biological variance and inferential variance is reported and estimated ([27]). It would be interesting to explore how RCD would work with such different types of error variance estimates.

In summary, whenever measurement error variances are available or can be predicted, our method RCD provides an opportunity to make reliable causal calls. This would immediately be of value to transform large-scale genetic and gene expression datasets into causal gene regulatory networks operating *in vivo* from yeast to human.

## Methods

### Background on MR framework

In the MR framework ([12]), a genetic variant denoted *L* (say a single-nucleotide polymorphism or SNP) is used as an instrumental variable to infer a causal relationship between two other trait variables denoted *G, T* (e.g., two gene expression traits in our setting) using mediation or other similar tests. More specifically, given an *L* that is correlated to both *G* and *T*, then under certain model assumptions pertaining to natural randomization of *L* that happens during meiosis and absence of confounding factors, statistical tests for correlation and conditional independence (also known as mediation) applied on the data collected on *L, G, T* can help distinguish between the causal models *L* → *G* → *T* and *L* → *T* → *G*, the (non-causal) independent model *G* ← *L* → *T*, and other similar models. Note that the non-causal independent model is called so, because *G* and *T* are independent when conditioned on *L*, but spuriously correlated otherwise (via the shared confounding factor *L*). Determining whether *L* is an instrument vs. a confounder is a key challenge in distinguishing between models where *G* and *T* are causally related (*G* regulating *T* or vice versa) vs. non-causally related.

Given that technical noise is inherent in any measurement including RNAseq measurements, we only have access to imprecise/noisy observations *G*′, *T* ′ of *G, T* respectively, and the natural question of interest then is: Can we develop a method that can work robustly even in the midst of different levels of technical noise in different genes?

### Background on CIT Causal Test

Since we extend the CIT ([5]) method, here we are summarizing the main points of the method. CIT is a mediation based causal discovery method which uses an instrumental variable (*L*) and checks whether its effect on the outcome variable (*T*) vanishes when conditioned on the mediating variable (*G*). The below description of the CIT is taken from ([5]). The three equations of the linear regression models are:

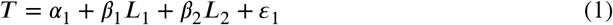

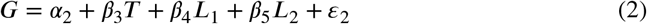

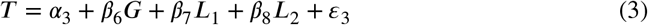

CIT tests causality conditions ([4]) using these 4 statistical tests:

1. *H*_0_ : {*β*_1_, *β*_2_} = 0, *H*_1_ : {*β*_1_, *β*_2_} ≠ 0; (*L* ∼ *T*),
2. *H*_0_ : {*β*_4_, *β*_5_} = 0, *H*_1_ : {*β*_4_, *β*_5_} ≠ 0; (*L* ∼ *G*| *T*),
3. *H*_0_ : *β*_6_ = 0, *H*_1_ : *β*_6_ ≠ 0; (*G* ∼ *T* |*L*),
4. *H*_0_ : {*β*_7_, *β*_8_} ≠ 0, *H*_1_ : {*β*_7_, *β*_8_} = 0; (*L* ⊥ *T* |*G*).

And among the four p-values, it takes the worst as the final p-value because the strength of all the tests is only as strong as the worst one.

In simulations and real-world data, CIT differentiates between the causal (*L* → *G* → *T*), reactive (*L* → *T* → *G*), and independent (*G* ← *L* → *T*) models using the cit.cp function implemented in its CRAN package ([28]) as follows. CIT first tests the *L* → *G* → *T* and *L* → *T* → *G* models, and then applies the *α* = 0.05 cutoff on the two resulting p-values as shown below to make the appropriate calls. The below description of model selection is taken from an earlier paper ([17]).

1. if p-value_*cit,G*→*T*_ < *α* and p-value_*cit,T* →*G*_ > *α*, CIT predicts causal model.
2. if p-value_*cit,G*→*T*_ > *α* and p-value_*cit,T* →*G*_ < *α*, CIT predicts reactive model.
3. if p-value_*cit,G*→*T*_ > *α* and p-value_*cit,T* →*G*_ > *α*, CIT predicts independent model.
4. if p-value_*cit,G*→*T*_ < *α* and p-value_*cit,T* →*G*_ < *α*, CIT predicts “No Call”.

### RCD Framework

#### Stage 1: Measurement Error Estimation

The aim of Stage 1 is to estimate the error variance 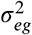, for any gene *G*. For cases when we have technical replicates, it is estimated directly as the sample variance of *G* across the replicates. If the technical replicates are unavailable, the error variance 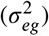 can be represented using different sources of technical variability, like sampling noise that arises due to differences in RNAseq library preparation steps ([15]) or noise due to alignment ambiguity ([27]) or can be any other unknown technical factor such as instrument error. Since Stage 2 of the framework is independent of any form of error variance estimates, in our current work we are modelling only sampling noise as the error variance estimate and show that we can obtain benefits even in this setup. As the mRNA fragments are selected randomly and independently from a large pool, they may or may not be sequenced and hence capturing it is a rare event that may reasonably be modelled as a Poisson distribution ([15, 20]). Since the count of technical replicates follows a Poisson distribution ([15]), we simulate dummy technical replicates to estimate the sampling or shot noise, the specifics of which are described next.

##### Estimating Noise in Normalized Gene Expression Data

A challenge in estimating measurement noise in normalized RNAseq data pertains to converting measurement noise in RNAseq read count space to noise magnitude in normalized RNAseq gene expression space (i.e., after standard normalization or log-transformation of the counts data). We are not aware of any analytical formula for this conversion, and we propose an empirical solution to address this challenge. We simulated technical replicate measurements of a dummy gene with the same average expression count as the original gene and subjected them to the same set of RNAseq normalization and log-transformation steps before estimating noise. In detail, let the observed count data be represented as an *n* × *m* matrix, where *n* is the number of genes and *m* is the number of samples. We first use a standard differential expression analysis method called DESeq ([20]) to estimate the sequencing depths or size factors *ŝ*_*j*_ for *j* = 1, …, *m* - these factors can be used to normalize the count data. The geometric mean of all size factors is used to estimate a single size factor *ŝ* that is robust to the differences in size factors across samples (note that geometric mean is considered a better average metric over arithmetic mean for gene counts data). For any given gene *gene*_*i*_, counts data for the dummy version of this gene, denoted *gene*_*ib*_, are simulated from a Poisson distribution with mean parameter 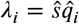, where 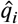 is the average of the DESeq-normalized-counts of *gene*_*i*_ across all samples ([20]). The simulated dummy count data is log-transformed with an offset of 0.5 to avoid issues with zero counts (specifically log_2_(*gene*_*ib*_ + 0.5)). The variance of this transformed dummy *gene*_*ib*_ is taken as the estimate of the noise variance. We bootstrap the above sampling process for 500 runs and average the estimated noise across these runs as a final estimate of the noise variance for *gene*_*i*_. The above process is repeated separately for each gene (i.e., *gene*_*i*_ for each *i*) to get gene-specific error variances 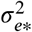. The process can be summarised as the below steps:

1. Take the count data matrix *k*_*ij*_ of size *n* × *m*, where *i* = 1, 2, …, *n* is the gene index and *j* = 1, 2, …, *m* is the sample index.
2. Estimate size factors: 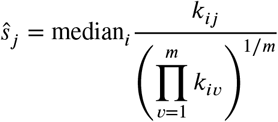
3. Normalize the count data matrix: 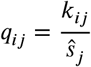
4. Estimate a single size factor: 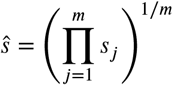
5. Repeat these two steps for *b* = 1, 2, …, *B*:
  a. Dummy gene counts: Simulate/draw *gene*_*ib*_ ∼ Poisson 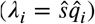, where 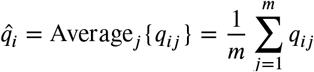. Here *gene*_*ib*_ refers to a vector of *m* independent draws from this Poisson distribution.
  b. Sample variance:

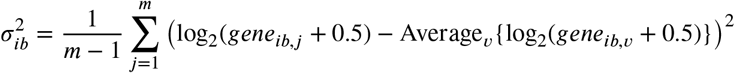
6. Noise estimate: 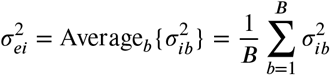

#### Stage 2: RCD Causal Tests

The aim of Stage 2 is to develop a causal discovery method that is robust against measurement errors by incorporating the noise estimates from Stage 1 (i.e., 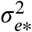 representing gene-specific error variances, with 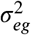 being the error variance of a particular gene *G*). In this section, we first describe our models explicitly in terms of their likelihoods and underlying assumptions such as linear causal relationships among variables; and next describe how the parameters of our linear models (with true variables) can be estimated from the corresponding noisy observed variables. These noise-corrected parameter estimates can then be used to adjust/correct four F-statistics based statistical tests of causality to account for noise (the same four tests proposed by CIT in a noise-free setting). Our techniques for incorporating noise magnitudes are similar to the errors-in-variables approach in linear regression ([29]), but we extend it to a causal discovery framework wherein noise magnitudes are incorporated to adjust not only parameter estimates of linear regression models but also associated residuals, F statistics and p-values pertaining to statistical tests of causality.

##### Model Assumptions, Description and Selection

In this work, we represent the causal relations among a trio of random variables (SNP *L* and two genes *G, T*) using appropriately defined linear regression (linear Gaussian) models. We consider three possible models as in CIT: causal (*L* → *G* → *T*), reactive (*L* → *T* → *G*), and independent (*G* ← *L* → *T*) models. We are asked to select one of these three models using noisy observations *G*′, *T* ′ of *G, T* respectively (the true gene expression values *G, T* are hidden from us), and noise-free measurements of *L*. We assume independent additive Gaussian noise distribution for the measurement errors of genes. The above model assumptions can be made more explicit by writing down the joint distribution (or likelihood of the model as a function of all model parameters *θ*). Consider the causal model above where *G* regulates *T*. Then, the joint distribution can be given by:

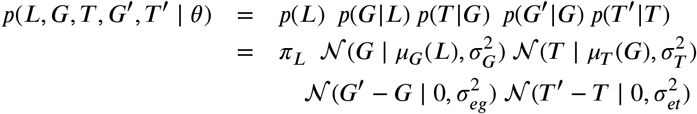

Here, *p*(·) denotes the probability density function (pdf) and 𝒩(*x*|*μ, σ*^2^) denotes the pdf of a Gaussian distribution with parameters *μ, <*^2^ evaluated at *x*. Note that *μ*_*G*_(*L*) above indicates that the expectation (average expression) of *G* is a function of *L* (specifically a linear function of *L* according to our model assumptions, as explained in detail below in the linear regression equations). Note that *π*_*L*_ is simply a parameter of the discrete or categorical distribution followed by *L*. Recall that 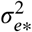 are the gene-specific error variances fixed in Stage 1 of RCD.

The joint distribution above can also be viewed as the joint distribution of a Bayesian network ([30]) comprising the directed edges: *L* → *G, G* → *T, G* → *G*^′^, and *T* → *T* ′. The description of the reactive and independent models would be similar, and can also be viewed as alternative Bayesian networks defined over the same set of three random variables.

Distinguishing between these three Bayesian network models to select one model using a series of association or conditional independence tests can then be viewed as structure learning of a Bayesian network using the constraints-based approach ([30]), which employs conditional independence tests to decide which edges to keep or remove in the learnt Bayesian network.

##### Coefficient Estimates from Noisy Data

The general linear regression equation is given as:

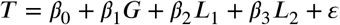

where *G* and *T* represent genes, and *L*_1_/*L*_2_ are variables encoding the genotype (or the number of non-reference alleles) as 0/0, 1/0 and 0/1 in that order. The ordinary least squares or maximum likelihood based estimates of the regression coefficients are given by:

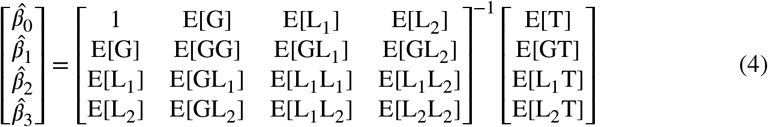

Here and in the rest of this paper, we assume that the sample averages are accurate approximations of model averages (i.e., expectations E[.]), which is a reasonable assumption if the sample size is sufficiently large. We also assume that the matrix that is being inverted in the above expression (call it *A*) is actually invertible. If that is not the case, we replace *A*^−1^ with the Moore-Penrose pseudo-inverse *A*^+^ to get a minimum-norm solution for *β*.

The above matrix formula can be reformulated in terms of observed noisy variables *G*′ and *T* ′ in our model described above, with *G*′ = *G* + *ε*_*eg*_, *T* ′ = *T* + *ε*_*et*_, and *ε*_*e*\*_ being the technical noise. We assume that the variance of the independent additive Gaussian measurement noise of a gene depends only on its average expression. Specifically, for any gene *X*,

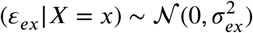

where 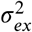 is the estimated error in Stage 1 of a dummy gene whose average expression is the same as^*e*^t^*x*^hat of the original gene *X* (in the count space, as detailed in Stage 1 of RCD framework). Note that this way of modeling *ε*_*ex*_ makes it independent not only of all other random variables in the system, but also of *X* (since the error *ε*_*ex*_ is *not* a function of the actual value x of *X*, but rather drawn independently based on the average expression count of *X*).

Hence, the estimates of various *G* and *T* statistics from *G*′ and *T* ′ are calculated as:

1. E[G] = E[G′]
2. 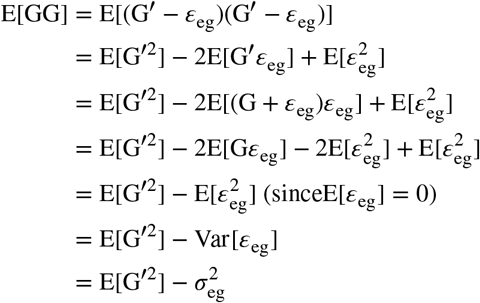
3. E[GL_1_] = E[(G′ − *ε*_eg_)L_1_] = E[G′L_1_]
4. E[T] = E[T′ − *ε*_et_] = E[T′]
5. 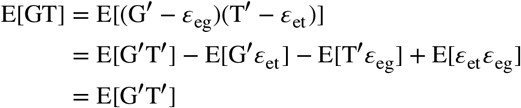

Using the above expressions, the reformulated estimates are given by:

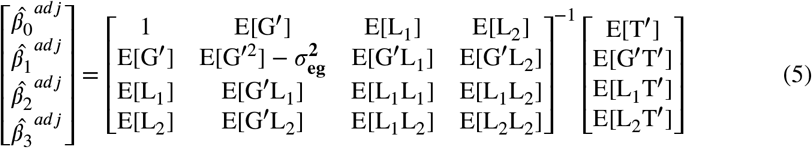

Please note that this reformulation of coefficient estimates offers one approach to correct measurement errors in the independent variables of a regression model, and suited our purpose in terms of performing reasonably well in simulated and real-world genomic data. However, other existing errors-in-variables regression modeling approaches, such as the total least squares methods or the Frisch scheme ([29]), could also be tried in the future.

##### Adjustment of F-statistic based Causality Tests to Handle Noise

The *F* statistic for testing the restriction of a complex linear regression model with *p* free parameters to a simple one with *q* free parameters is given by ([31]):

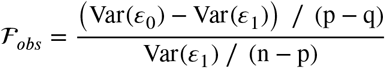

Here, Var(*ε*_0_) and Var(*ε*_1_) are the variance of residual of the nested simple and complex model respectively. This statistic follows an *F*_*p*−*q,n*−*p*_ distribution under the null hypothesis (*H*_0_) that the simple model is the true model. Under the alternate hypothesis (*H*_1_), the complex model is the true model generating the data.

We next show how RCD incorporates measurement error estimates into the four statistical tests of CIT. In the first three tests, *H*_0_ is given by the simple model and *H*_1_ by the complex model as above; but the fourth test is the converse as detailed below.

1. Is *L* and *T* associated: (*L* ∼ *T*)?

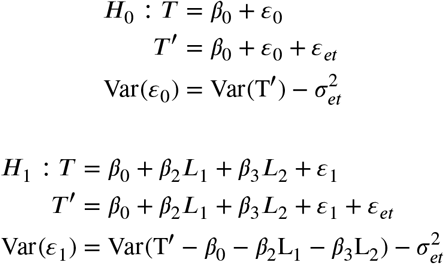

Since the variables in the LHS (Left Hand Side) of the above regression equations are noisy and all the RHS (Right Hand Side) variables are noise-free, we can use a formula similar to Formula 4 to estimate the above coefficients.
2. Is *G* and *T* associated given *L*: (*G* ∼ *T*| *L*)? **Adjusted F statistic Formula**

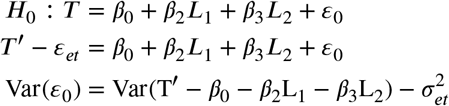

Since only the LHS variables are noisy, we use a formula similar to Formula 4 to estimate the above coefficients.

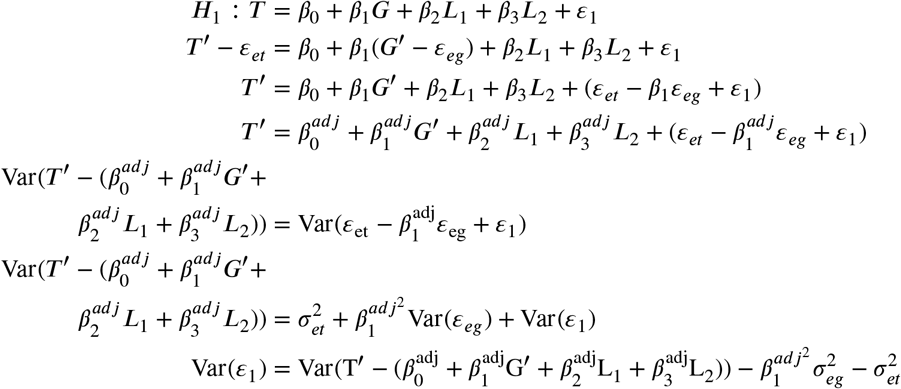

To handle noise in both LHS and RHS variables, we use adjusted Formula 5 to estimate the above coefficients.
3. Is *L* and *G* associated given *T* : (*L* ∼ *G* |*T*)? **Adjusted F statistic Formula**

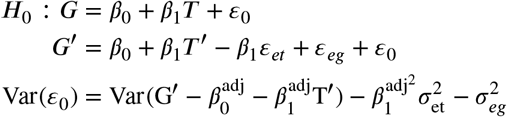

We use a formula similar to adjusted Formula 5 to estimate the above coefficients.

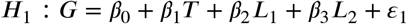

Similar to test 2, by interchanging *T* and *G* we obtain the result as follows:

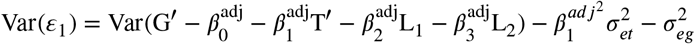

We use adjusted Formula 5 (after interchanging *T* and *G*) to estimate the above coefficients. **Empirical Null Distribution using the Bootstrap and *F* * statistic**: For the test (*L* ∼ *G*|*T*), the null model is *L G*|*T* ⊥. We followed an empirical approach to obtain *F*\* distribution under the null. We use the below approach to simulate from *L* ⊥ *G*|*T* in noisy settings.

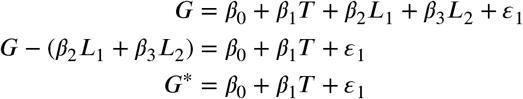

where *G*^*^ = *G* − *β*_2_*L*_1_ − *β*_3_*L*_2_. From this we can see that dependency between *G* and *L* is broken while maintaining other dependencies, i.e. (⊥ ^*^), but when we have a noisy version of *G* as *G*^′^, we can get a similar result as follows:

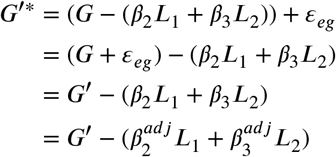

Since (*L* ⊥ *G*′*|*T* ′), we can compute *F* * statistic under null distribution as follows:

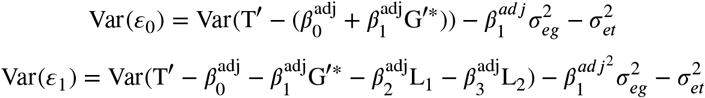

We use formula similar to adjusted Formula 5 to estimate the above coefficients. We can bootstrap from this (*L, G*′*, *T* ′*) dataset *B* times (for a sufficiently large *B*) to get the empirical distribution of *F* *, and compare the observed test statistic *F* directly against this empirical distribution of *F* * to get the relevant p-value, as done in CIT ([5]).
4. Is *L* and *T* independent given *G* : (*L* ⊥ *T* |*G*)?

Here we use an equivalence test where the alternate hypothesis is (conditional) independence rather than association; so in contrast to the above three tests, the complex model corresponds to the null hypothesis *H*_0_, and the nested simple model to *H*_1_.

##### Adjusted F statistic Formula

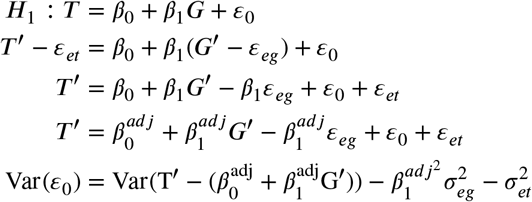

We use a formula similar to adjusted Formula 5 to estimate the above coefficients.

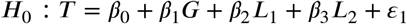

It is the same as test 2 (*H*_1_), so we obtain the below result:

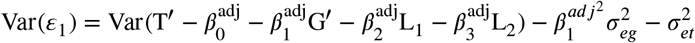

We use adjusted Formula 5 to estimate the above coefficients.

##### Empirical Null Distribution using the Bootstrap and *F* * Statistic

For the test (*L* ⊥ *T* |*G*), the null model is *G* ← *L* → *T*. We followed an empirical approach to obtain *F* * distribution under the null. We use the below approach to simulate from *G* ← *L* → *T* in noisy settings. A random variable *G*′* is simulated such that *L* and *G*′* is associated, but dependence between *G*′* and *T* is broken. We do so, as in CIT ([5]), using these steps:

a. First, we estimate the association parameters by regressing *G*′ on *L* as

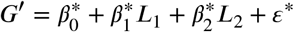

and using a formula similar to Formula 4.
b. Then, the residual vector *ε*\* is randomly permuted and used along with the estimated parameters to obtain *G*′* as

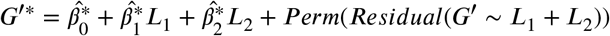

Now, the *F* * is estimated as follows:

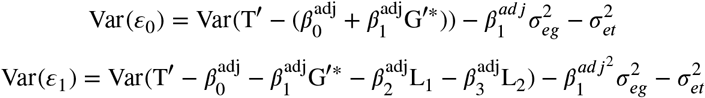

We use formula similar to adjusted Formula 5 to estimate the above coefficients. We bootstrap *B* times (for sufficiently large *B*), i.e., repeat the (independent) random permutation step above *B* times to obtain an empirical distribution of *F* *. Against this empirical null distribution, the observed test statistic *F* is compared to obtain a p-value, again as in CIT.

### Simulation Setup

Simulations were performed on different parameter combinations. *L* is the instrument variable, *G* is the hidden causal mediator variable, *T* is the hidden outcome variable, and *G*′ and *T* ′ are the corresponding observed noisy variables as shown in Figure 1. The simulation settings are taken from ([17]). Causal data is simulated following the model as described:

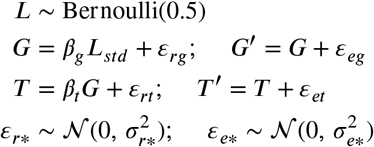

*L*_*std*_ above refers to the standardized (z-score transformed) *L*. Residual variances (i.e., variances unexplained by the causal factor) are captured using independent Gaussian-distributed random variables *ε*_*r*\*_, whereas measurement error variances are captured using the independent Gaussian variables *ε*_*e*\*_. In the expressions above and in the text, * is a shorthand for *G, T*, which indicates that the corresponding expression applies separately for each of the genes *G* and *T*.

We can set the *β*_*_ and 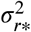 parameters in such a way that:

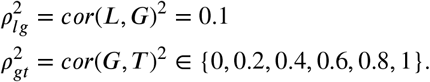

We specifically set *β*_*g*_ = *ρ*_*lg*_, 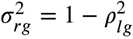, and *β*_*t*_ = *ρ*_*gt*_, 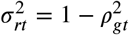. The range of values for parameters above, along with the range of values of the noise magnitudes and sample size *n* below, results in a total of 648 configurations for the causal model. For each configuration, CIT and RCD are run 100 times.

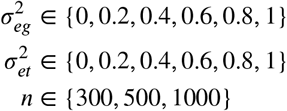

Note that the above configurations correspond to the causal model *L* → *G* → *T* (including the model where *G, T* link is severed due to *cor*(*G, T*)^2^ = 0). For the non-causal or independence model *G* ← *L* → *T*, we simulate data by changing the dependence of *T* from *G* to *L*, i.e., by replacing the *T* expression in the causal model above by *T* = *β*_*lt*_*L*_*std*_ + *ε*_*rt*_, with *β*_*lt*_ set to *ρ*_*lt*_ and 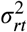 set to 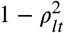, and leaving other models/expressions unchanged.

We used the implementation of CIT in the CRAN package ([28]). We implemented RCD using the R programming language ([32]).

### Evaluation Setup for Yeast GRN Inference

#### Estimating Measurement Noise

To estimate shot noise variance, we applied the Stage 1 procedure (see Methods section on “RCD Framework”) to the 1012-segregants yeast mRNA counts data^1^.

##### Selecting Strongest cis-eQTLs

To run CIT or RCD, we need an eQTL which can be used as the *L* in the model. We used cis-eQTLs and cis-egene reported in ([33]). For a single cis-egene, there can be more than one eQTLs; so we grouped eQTLs based on cis-egene and selected the strongest cis-eQTL for each cis-egene based on the absolute correlation coefficient. We obtained a total of 2433 eQTLs, out of which 2044 are associated with one cis-egene, and 337, 44, 6, and 2 are associated respectively with two, three, four and five cis-egenes.

#### Target Network Inference

The mRNA counts matrix is available for 5720 genes across 1012 yeast segregants. We applied DESeq (size-factors-based) normalization on this counts data, followed by the log-transformation, log_2_(DESeq_normalised + 0.5). Measured covariates provided in ([33]) are regressed out from the log-transformed data using categorical regression model and causality analysis is done on the final regressed-out gene expression data. From YEASTTRACT+ database ([24]), DNA plus Expression binary regulation matrix on previously studied 80 TFs and 3394 TGs ([34]) is used as ground-truth network, where for each TF/Gene pair, the Regulatory Association (RA) is represented by 0 or 1, representing a non-existing or existing association, respectively. Excluding self-regulations, it has 62052 existing and 207933 non-existing associations. A subset of this complete list of trios involving genes with high measurement error is also used in our analysis. A trio is called a noisy or high measurement error trio with respect to a certain cutoff, for instance 40%, if: 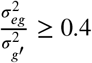 or 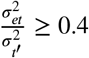. Here the total gene variance in the denominator (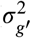 or 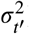) is the sample variance of the gene after adjustment for covariates (specifically two covariates in the case of our yeast dataset, optical density indicating growth-rate and batch/date of RNAseq processing of the sample).

### Dataset Description for Human Tissue

We performed skeletal muscle analysis on the NIH GTEx ([25]) V7 data (dbGaP Accession phs000424.v7.p2).

#### Estimating Measurement Noise

GTEx has provided mRNA counts data for all tissues in data-source ^2^. Using the annotation data-source^3^, we keep only samples specific to skeletal muscle, and then estimate shot noise variance by applying our RCD framework’s Stage 1 procedure described above to the skeletal muscle mRNA counts data.

#### Selection of trios

To get a query list of trios (*L, G, T*) on which to run RCD, we considered *L* and *G* directly given in the GTEx portal as cis-eQTL data-source^4^. For multiple eQTLs of the same cis-egene, we used only one based on the best q-value. We used a *q* ≤ 0.05 cutoff as recommended in the GTEx portal to get significant cis-eQTLs. This gives us a list of (*L, G*), where *L* is the cis-eQTL of the corresponding cis-egene *G*. Corresponding to each significantly identified eQTL association (*L, G*), we used significant trans-egenes identified by Matrix eQTL ([35]) that are also at least 1 Mb distance away from *L* as possible *T* genes. Pairs (*L, T*) with trans-association p-value < 10^−5^ is considered as significant, which gives a query list of trios (*L, G, T*) for downstream analysis. Covariates and other PEER factors given in the GTEx data-source^5^ are used to adjust the data before causality analysis.

## Acknowledgments

We thank members of our BIRDS (Bioinformatics and Integrative Data Science) research group for their valuable input during the presentations of this work. The research presented in this work was supported by Wellcome Trust/DBT grant IA/I/17/2/503323 awarded to MN.

## Supplementary Material

**Supplementary Figure S1.**
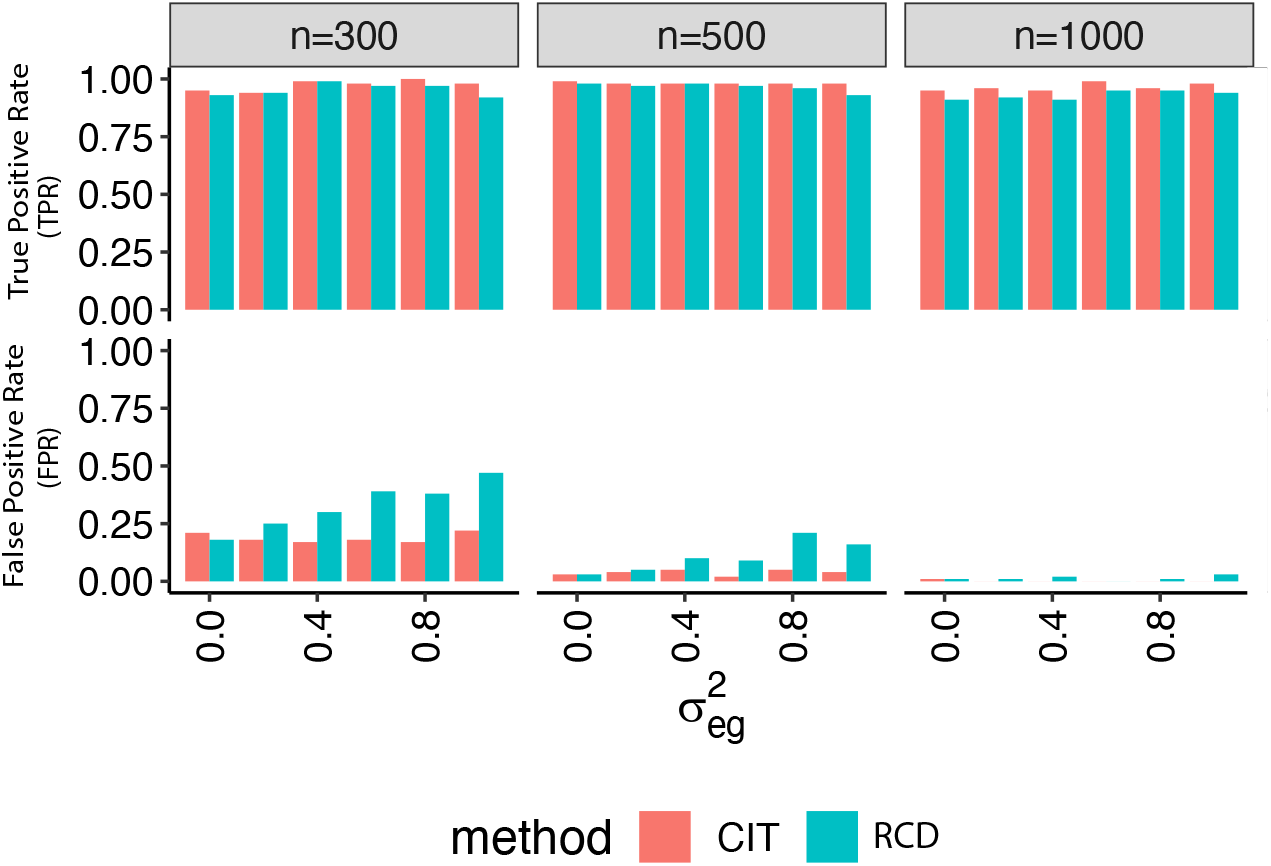
Performance comparisons of RCD on simulated independent vs. causal pairs. (top) Top graph plots the true positive rate (fraction of all true independent relations that are inferred as independent by RCD or the baseline method CIT), when data pertaining to 100 true independent pairs are simulated according to the independent model *G*← *L* → *T* (with correlation coefficient *ρ*_*gt*_ = 0 004; note that this is spurious correlation due to *L*). (bottom) Bottom graph plots the false positive rate (fraction of all causal pairs inferred as independent by a method), when data on 100 causal pairs are simulated using the model *L* → *G* → *T*. Measurement error in mediator 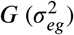 is varied along the x-axis keeping the measurement error in trait *T* fixed at 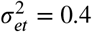. Power (TPR) of both the methods are similar, whereas RCD exhibits somewhat higher^*et*^FPR than CIT at low sample sizes (column panels show different sample sizes: *n* = 300, 500, 1000).

**Supplementary Figure S2.**
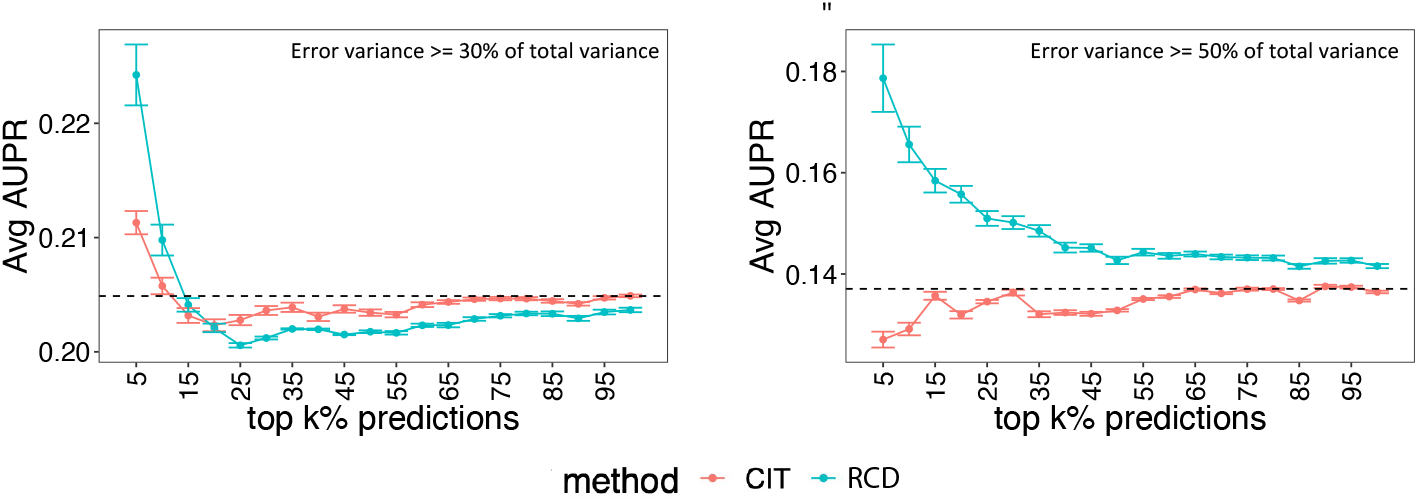
Performance of our RCD relative to the baseline CIT on yeast dataset at different noise cutoffs. The goal is to recover ground-truth causal regulation of TFs→TGs, after keeping only those interactions for which either TF or TG have high measurement errors. Performance on ground-truth causal interactions with TF or TG having error variance at least 30% of the total variance (left) and at least 50% of the total variance (right).

**Supplementary Figure S3.**
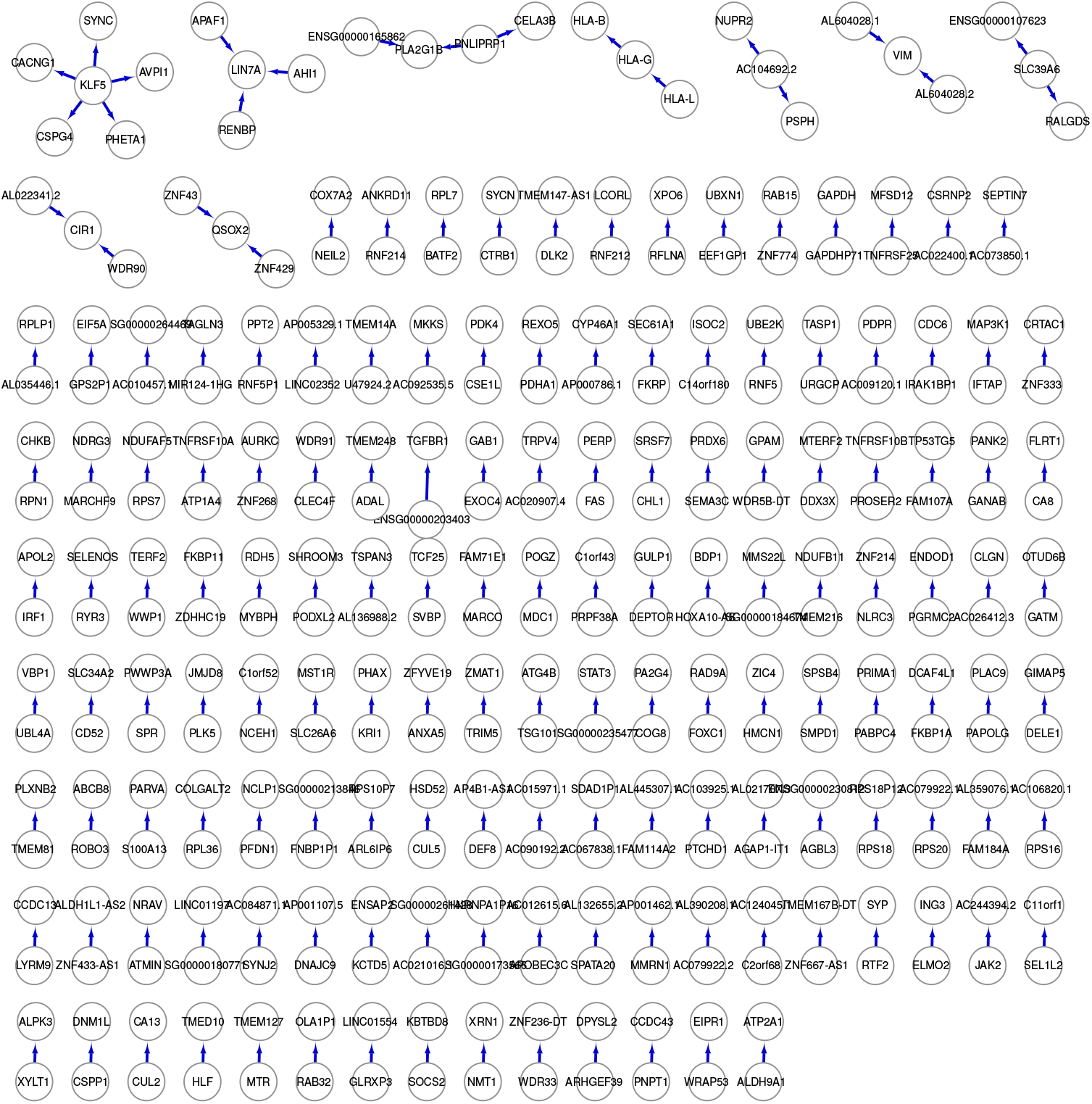
Entire human muscle gene regulatory network discovered by RCD. A human muscle gene regulatory network comprising 314 genes and 164 directed edges inferred by RCD from NIH GTEx skeletal muscle tissue data. Direction *G* → *T* is predicted if p-value_*RCD,G*→*T*_ < 0.05 and p-value_*RCD,T* →*G*_ > 0.05.

**Supplementary Figure S4.**
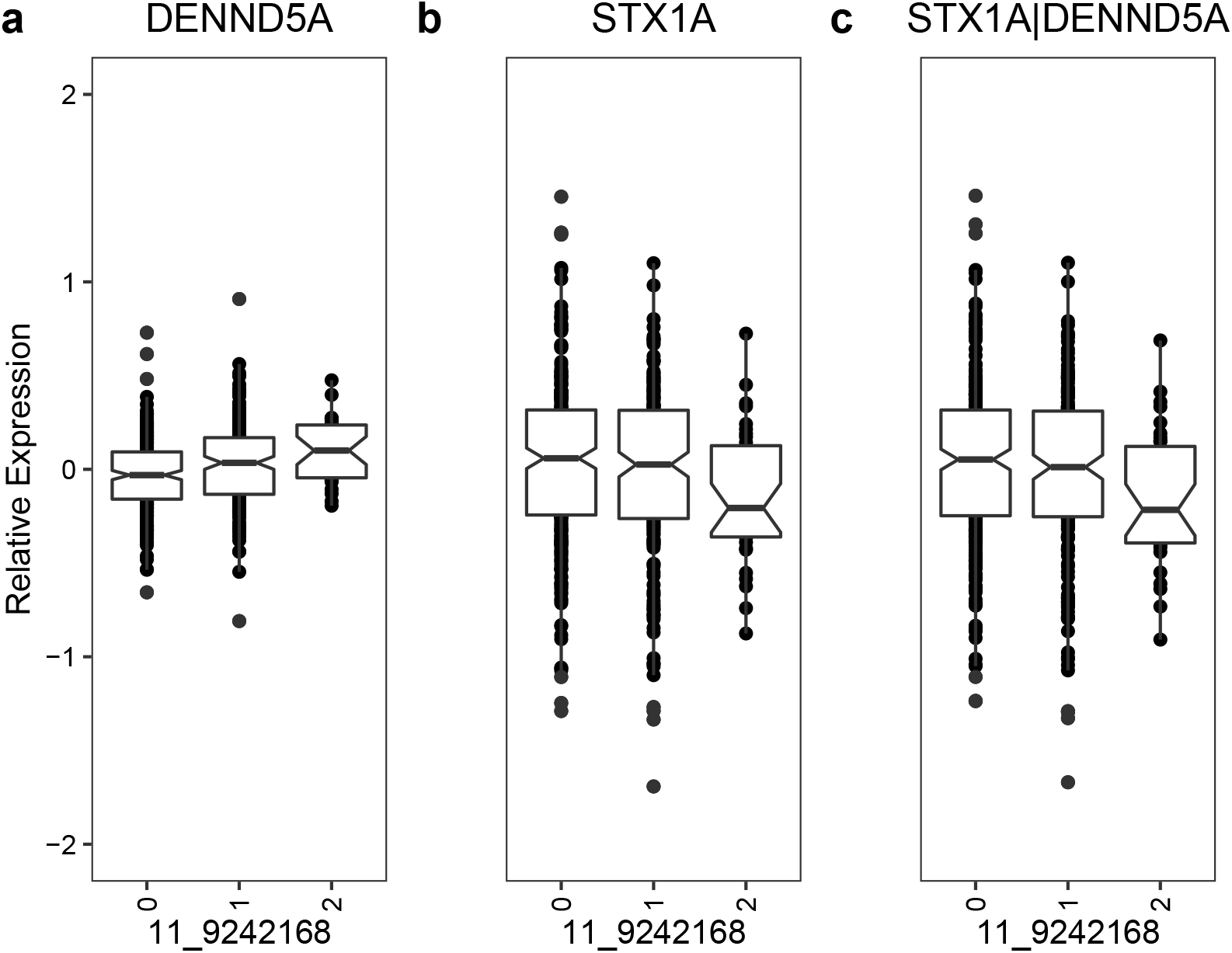
Illustration of how RCD detects spurious correlation between two genes. An illustration of two genes that RCD revealed to be spuriously coexpressed due to the effect of a shared confounding SNP. A SNP *L* at 11_9242168, encoded as 0, 1 or 2 based on the number of copies of the non-reference allele, is correlated to both genes *G* and *T* as shown in (a) and (b) respectively. An independent model, *G* ← *L* → *T* is supported by the data as (a) *L* → *G*: SNP is associated with a candidate regulator gene, DENND5A (b) *L* → *T* : SNP is associated with a candidate target gene, STX1A and (c) importantly, *L*/⊥*T*|*G*: effect of SNP on STX1A does not vanish once conditioned on the candidate regulator DENND5A.

## Footnotes

1 Yeast mRNA counts data https://github.com/joshsbloom/eQTL_BYxRM/blob/master/RData/counts.RData

2 GTEx_Analysis_2016-01-15_v7_RNASeQCv1.1.8_gene_reads.gct.rds https://www.gtexportal.org/home/datasets

3 GTEx_v7_Annotations_SampleAttributesDS.txt https://www.gtexportal.org/home/datasets

4 Muscle_Skeletal.v7.egenes.txt.gz https://www.gtexportal.org/home/datasets

5 GTEx_Analysis_v7_eQTL_covariates https://www.gtexportal.org/home/datasets

